# Mettl15-Mettl17 modulates the transition from early to late pre-mitoribosome

**DOI:** 10.1101/2024.12.18.629302

**Authors:** Yury Zgadzay, Claudio Mirabello, George Wanes, Tomáš Pánek, Prashant Chauhan, Björn Nystedt, Alena Zíková, Paul C. Whitford, Ondřej Gahura, Alexey Amunts

## Abstract

The assembly of the mitoribosomal small subunit involves folding and modification of rRNA, and its association with mitoribosomal proteins. This process is assisted by a dynamic network of assembly factors. Conserved methyltransferases Mettl15 and Mettl17 act on the solvent-exposed surface of rRNA. Binding of Mettl17 is associated with the early assembly stage, whereas Mettl15 is involved in the late stage, but the mechanism of transition between the two was unclear. Here, we integrate structural data from *Trypanosoma brucei* with mammalian homologs and molecular dynamics simulations. We reveal how the interplay of Mettl15 and Mettl17 in intermediate steps links the distinct stages of small subunit assembly. The analysis suggests a model wherein Mettl17 acts as a platform for Mettl15 recruitment. Subsequent release of Mettl17 allows a conformational change of Mettl15 for substrate recognition. Upon methylation, Mettl15 adopts a loosely bound state which ultimately leads to its replacement by initiation factors, concluding the assembly. Together, our results indicate that assembly factors Mettl15 and Mettl17 cooperate to regulate the biogenesis process, and present a structural data resource for understanding molecular adaptations of assembly factors in mitoribosome.

---

In mitochondria, messenger RNA (mRNA) translation and protein synthesis are performed by the mitoribosome in association with the regulatory complex LRPPRC-SLIRP^1^ and the OXA1L insertase at the inner mitochondrial membrane^2,3^. The mammalian mitoribosome consists of three mitochondria-encoded RNA molecules with 19 modified nucleotides and at least 82 nuclear-encoded proteins^4-6^. The formation of this complex machinery involves progressive assembly through the recruitment of assembly factors that act primarily on the ribosomal RNA (rRNA), triggering its gradual folding and modification, while mitoribosomal protein modules are formed^7-9^. This allows for productive maturation through defined states that ultimately leads to the catalytic mitoribosome^10^. Perturbations in the assembly pathway can underlie association of mitoribosomal dysfunction with various diseases^11-14^.

Structural studies on *Trypanosoma brucei* mitoribosomal complexes showed that it is a good model for understanding fundamental principles of mitoribosomal assembly because its native pre-mitoribosomal complexes are biochemically more stable and contain most of the assembly factors observed in mammals and other eukaryotes^15-19^. For example, *T. brucei* mitoribosomal large subunit biogenesis involves at least seven assembly factors shared with humans, including GTPases GTPBP7, MTG1, pseudouridinase RPUSD4, and methyltransferase MRM. The structures of intermediates with these factors allowed for a better understanding of their roles in the mitoribosomal assembly pathway^17,18^.

The mammalian mitoribosomal small subunit (mtSSU) is highly dynamic and contains 12S rRNA with nicotinamide adenine dinucleotide associated with an rRNA insertion, 30 mitoribosomal proteins, and two iron–sulfur (Fe-S) clusters^20-23^. The structure is arranged into two main regions defined as the body and head. The latter binds LRPPRC-SLIRP, which regulates mRNA delivery to a dedicated channel during translation initiation and undergoes conformational changes to accompany the movement of mRNA during translation cycle^4,5,24^. In skeletal muscle, exercise training-induced signalling leads to enhanced mitoribosomal activity that can bypass LRPPRC-SLIRP^25^. The assembly path involves at least 11 factors that facilitate binding of mitoribosomal proteins and construct the solvent-exposed surface of the rRNA, including the mRNA channel and the decoding centre at the interface between the body and head^8^.

Structural studies have revealed that stable assembly intermediates of the small subunit can be divided into ‘early’^26^ and ‘late’^22^ stages, each relying on distinct methyltransferases, Mettl17 and Mettl15, respectively. Mettl17 is also an Fe-S binding protein that serves as a checkpoint for mitochondrial translation^27^. However, the transition from the early to late stage, including the interaction between Mettl15 and Mettl17, has never been observed. It is also not clear what drives the release of Mettl17, which promotes maturation, primarily due to limitations in the experimental design.

Studying mitoribosomal assembly intermediates experimentally is challenging because interactions between assembly factors are often dynamic, and the transient states can undergo dissociation when isolated for structural analysis. In addition, adding tags to assembly factors for protein purification can interfere with native interactions, and knockout strains might exhibit non-productive off-path configurations of pre-mitoribosome, thus compromising the interpretation. However, the recent development of new computational tools for the analysis of protein-protein interactions^28,29^ enabled studies on large nucleoprotein complexes involved in gene expressions and associated with transient modifying enzymes^23,24^. Thus, *in silico* approaches can reveal direct interacting partners and propose models of sequential assembly of macromolecular complexes.

Here, we used the cryo-EM map of *T. brucei* mtSSU assembly intermediate^15^ and structural models of human early^26^ and late^22^ intermediates. Leveraging recent computational advancements, we performed *AlphaFold2*^30^ analysis and molecular dynamics simulations, to generate *in silico* models for previously undescribed states. This approach enabled us to propose a sequential mechanism that explains the structural basis for the Mettl15-Mettl17 function on the pre-mitoribosome. PyMol^65^ sessions are available for all the states as a Resource (Supplementary Information).

## RESULTS

### Mapping unassigned regions in the pre-mitoribosome uncovers Mettl15-Mettl17 heterodimer

A mtSSU intermediate from *T. brucei* has been previously studied by cryo-EM, but several regions in the map remained unassigned^15^. Using the data from recently published structures of mammalian pre-mitoribosomal intermediates^22,26^, we analysed the *T. brucei* maps and identified a number of previously undescribed structural elements (Table S1).

First, we detected homologs of RbfA and Mettl15 (previously referred to as mt-SAF18 and mt-SAF14, respectively), both of which are associated with Mettl17 (mt-SAF1) (Figure 1A). RbfA is a KH-domain containing assembly factor (Figure 1B) that scaffolds decoding center rRNA elements, contacts the 3’end of rRNA, and occupies the mRNA channel during ribosomal assembly in bacteria and mitochondria^22,31-34^. Mettl15 is a class I SAM-dependent N4-methylcytidine (m^4^C) methyltransferase of bacterial origin that modifies the mtSSU rRNA at position C1486 (human numbering)^35-38^. Mettl17, in contrast, is a putative methyltransferase with no specific target^39,40^, and the disruption of its interaction with the pre-mitoribosome impairs other methyltransferases as well^35,36,38^. Structurally, RbfA is anchored to the complex by its N-terminal extension, with the C-terminal domain binding Mettl17 and the C-terminal extension binding Mettl15. Together, these elements stabilize the subcomplex in a way that Mettl15 and Mettl17 form a heterodimer that is bound in the cleft between the head and body (Figure 1C). The Mettl15-Mettl17 heterodimer has the Complexation Significance Score of 0.695, this score is defined as the maximal fraction of the total free energy of binding^62^, which indicates a specific interface, and the interaction surface area is 4380 Å^2^. In total there are 43 hydrogen bonds and ten salt bridges that stabilize the Mettl15-Mettl17 heterodimer (Table S1), which contribute to two main interfaces. The first interface comprises the N-terminal part of Mettl17 (residues 45-95) and C-terminal part of Mettl15 (residues 402-470). The second interface involves catalytic domains of both Mettl17 and Mettl15 (residues 484-512 and 176-198, respectively). These data suggest that in *T. brucei* Mettl15 and Mettl17 form a stable complex, which has not been observed in other species.

**Figure 1.**
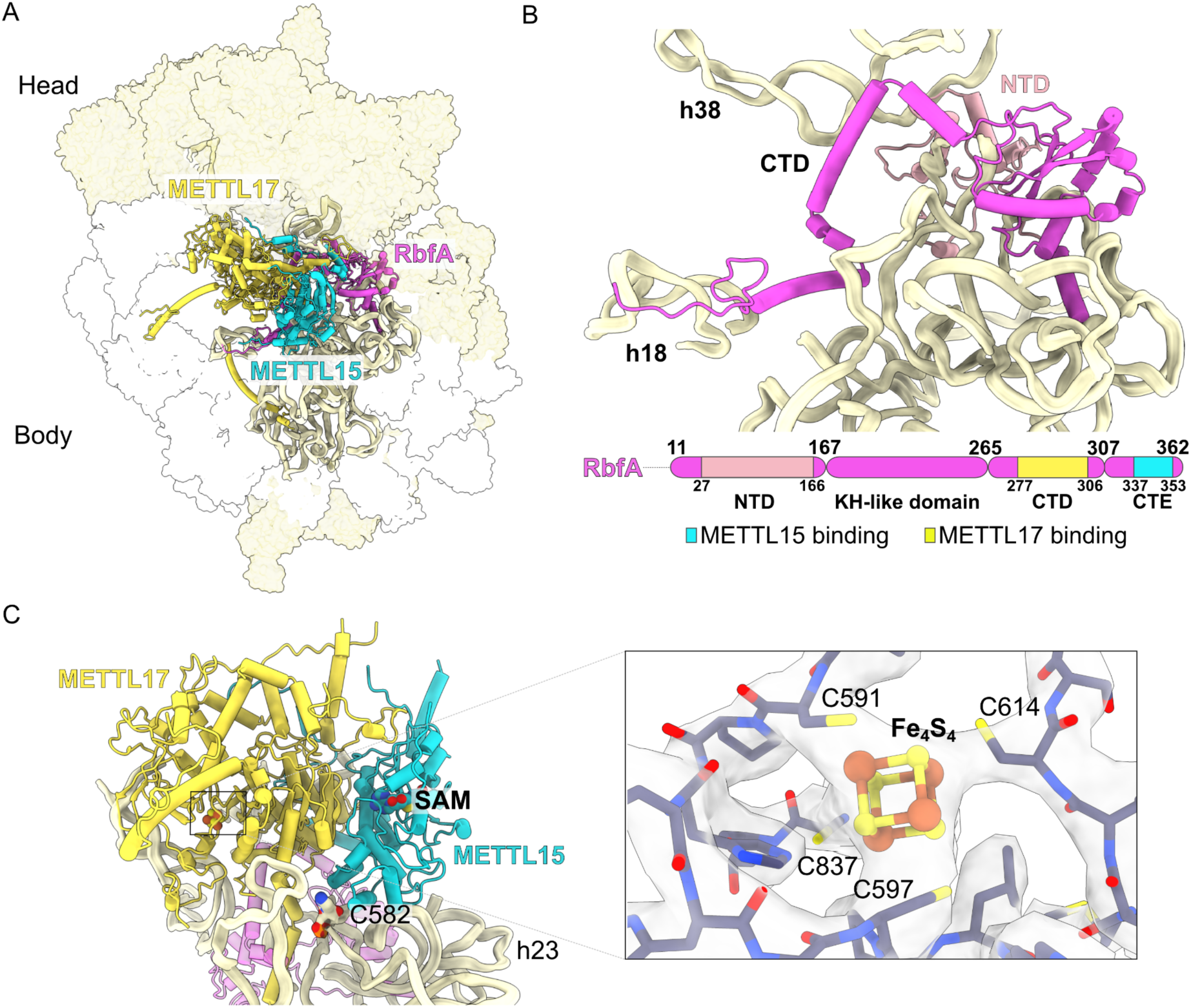
Dimer of Mettl15 and Mettl17 is found with RbfA in a *T. brucei* pre-mtSSU. **(A)** Structure of pre-mtSSU highlights Mettl15 (yellow), Mettl17 (cyan), RbfA (magenta) and rRNA (pale-yellow ribbon). Other assembly factors and mitoribosomal proteins are shown as white and yellow surfaces, respectively. **(B)** Structure and schematic representation of RbfA and its interaction with rRNA. Mettl15 and Mettl17 interacting regions are indicated. **(C)** Structure of the Mettl15-Mettl17 heterodimer with cofactors. A close-up view shows Fe_4_S_4_ with its coordinating residues.

Both methyltransferases in the structure contain a functional prosthetic group S-adenosyl methionine (SAM) (Figure 1C). In Mettl15, SAM is located 42 Å away from the methyltransferase target residue cytidine 582 (C582, equivalent of human C1486), implying a non-catalytic conformation. The Mettl15 conformation is different from that observed in the mammalian m^4^C1486-containing post-catalytic precursor^22^, thus implying a pre-catalytic state. In Mettl17, there is a density corresponding to an iron-sulphur Fe_4_S_4_ cluster, consistent with the mammalian^26^ and yeast^27^ homologs. Thus, the previously proposed role of Mettl17 as an oxidative stress sensor and an Fe-S checkpoint for mitochondrial translation^27,41^ may be conserved in a broad range of eukaryotes.

In addition, we assigned an uninterpreted region of density to trypanosomal assembly factor mt-SAF38 (Figure S1A). Its overall fold is similar to a thioesterase, expanding a list of enzyme homologs identified in mitoribosomal subunits or their precursors^42^. Finally, we identified 13 hammerhead-shaped densities ranging from 17 to 22 Å in length, coordinated by tryptophan residues within a helix-loop-helix motif of pentatricopeptide repeat (PPR) proteins, which likely represent cofactors such as acetyl coenzyme A (acetyl-CoA) (Figure S1B).

### Evolutionary conservation of Mettl17 suggests its role in recruiting Mettl15

To determine whether the Mettl15-Mettl17 heterodimer is a group-specific feature or may be widespread, we searched for these two methyltransferases in genomes of diverse eukaryotic organisms, followed by phylogenetic analysis. The search identified Mettl17 in 126 out of 134 organisms covering all major eukaryotic lineages. The cysteine residues coordinating the Fe_4_S_4_ cluster in mammals and trypanosomes are conserved in most identified Mettl17 homologs. While Mettl15 is present in fewer organisms, it was identified in nearly all species where Mettl17 was present (Figure 2, Supplementary Data 1&2, Figures S1, S2). This suggests that Mettl17 may be a prerequisite for the incorporation of Mettl15 into the pre-mitoribosome. Mettl17 is essential for mitochondrial translation in human cells^39^, for mitoribosomal assembly, translation and viability in *T. brucei*^15,43^, and for respiration in budding yeast^44^, but there is currently no evidence for the methyltransferase activity of this protein in any organism. Instead, human Mettl17 is required for methylation of rRNA by Mettl15^39^. Thus, consistently with other enzymes that adopted a structural role in the mitoribosome^45^, Mettl17 is an essential and conserved protein with no specific methylation target, whose primary function may be to facilitate Mettl15 integration into the pre-mitoribosome.

**Figure 2.**
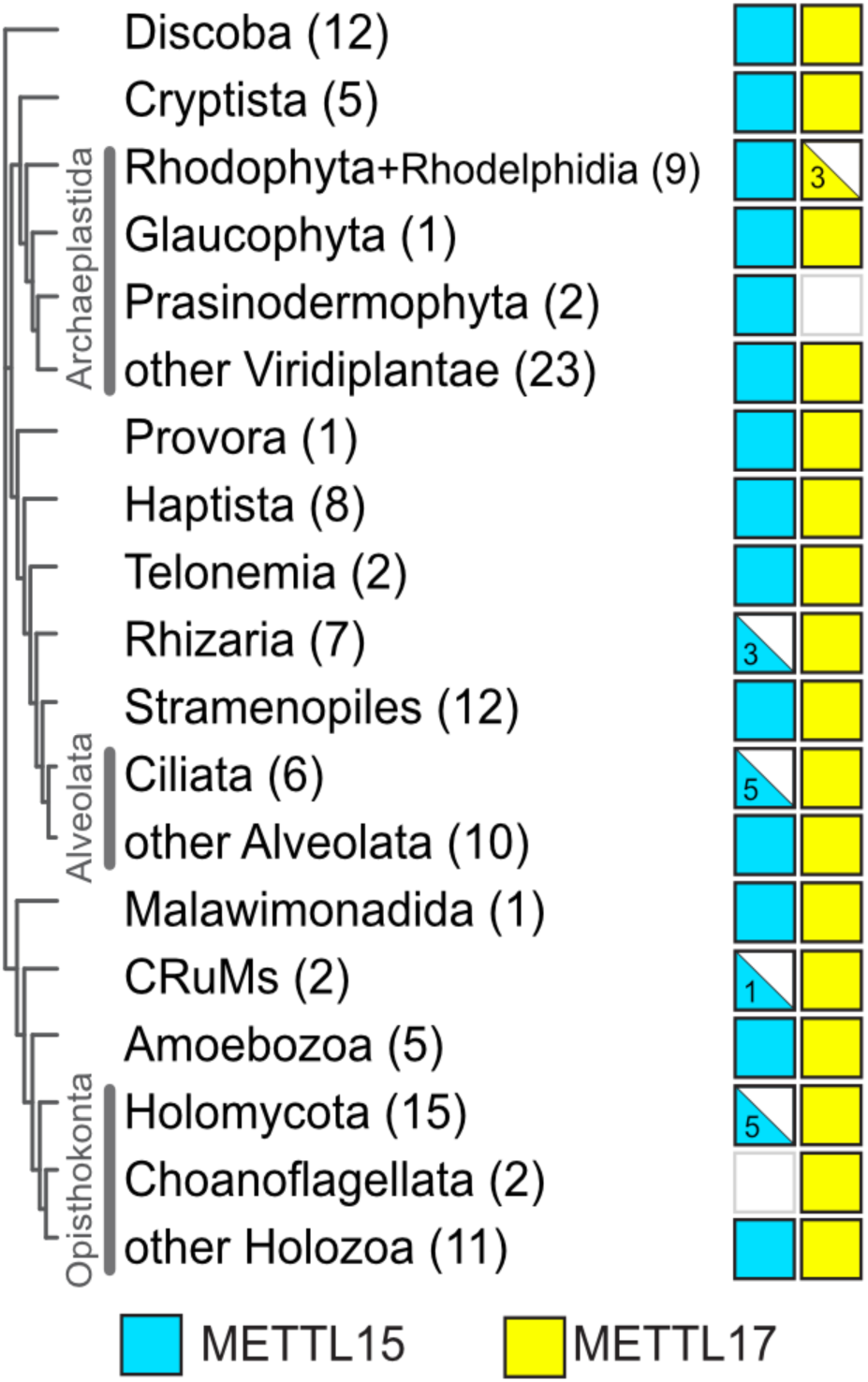
Mettl17 and Mettl15 are widely distributed across eukaryotes. Distribution in major eukaryotic groups mapped onto a phylogenetic tree^62^. The numbers in parentheses indicate the number of organisms searched in the respective group. Filled symbols represent presence in all organisms, half-filled symbols indicate presence in a subset of organisms, and empty symbols indicate absence.

### Mettl15 associates with Mettl17 on the pre-mitoribosome during early assembly stage

To clarify at which stage Mettl15 associates with Mettl17 on the pre-mitoribosome, we used structural models of human early^26^ and late^22^ intermediate as references. The early intermediate contains Mettl17 and another methyltransferase, TFB1M (PDB ID 8CSP), whereas the late stage contains Mettl15 in a different conformation (PDB ID 7PNX). We generated *AlphaFold2* (AF2)^28^ models of human Mettl17-TFB1M and Mettl15-TFB1M (Figure 3A,B). The two models obtained similar protein interface (ipTM) scores of 0.66 and 0.59, respectively, which would indicate reasonable confidence, according to the most recent benchmarking of AF prediction of multi-chain protein complexes^46^. The AF2 model of Mettl17-TFB1M corresponds to the experimental dimer of the two proteins in the cryo-EM structure of the early state^26^, supporting the computational approach. We then used TFB1M as an anchoring point for superposition of Mettl15-TFB1M from the predicted model onto the early intermediate (Figure 3C). The superposition shows that Mettl15 is compatible with Mettl17, except minor clashes observed between a loop in Mettl17 (residues 220-247) and Mettl15 (residues 205-249). However, the Mettl17 loop has relatively high B-factor compared to rigid parts of the protein in current structures (Figure S2), indicating it is rather flexible and could attain alternative conformations when in complex with Mettl15. This suggests that Mettl15 could be structurally co-localized with Mettl17, TFB1M and RbfA on the pre-mitoribosome. This is further supported by biochemical evidence, as TFB1M readily co-immunoprecipitate with Mettl15^36^. Thus, Mettl15 potentially associates with the pre-mitoribosome during the early assembly stage, possibly co-constituting a state with all three methyltransferases bound (Figure 3C).

**Figure 3:**
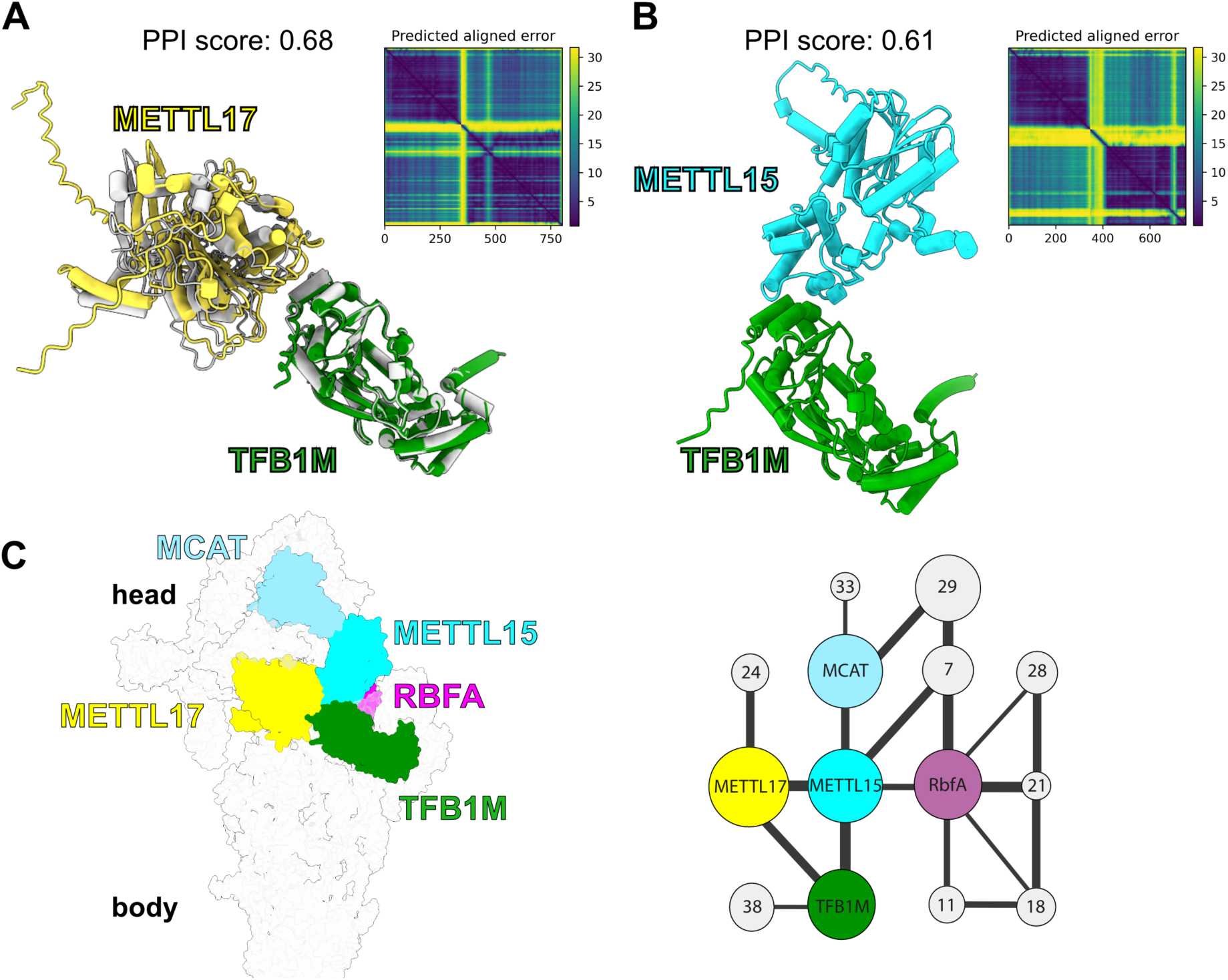
Modelling of early pre-mtSSU with Mettl15. **(A)** AlphaFold model of Mettl17-TFB1M superposed with the experimentally determined model of the early state (grey, PDB 8CSP). **(B)** AlphaFold model of Mettl15-TFB1M. **(C)** Left, model of early pre-mtSSU with all three methyltransferases, including Mettl15. Right, schematics of protein-protein interactions of methyltransferases, other assembly factors (colored nodes), and mitoribosomal proteins (grey nodes). The node size corresponds to relative molecular mass of protein subunits, and the connector width corresponds to the relative solvent accessible interface area buried between the subunits, calculated with PDBePISA v.1.52^63^.

### Pre-mitoribosome with Mettl15 and Mettl17 represents a pre-catalytic state

To establish the context of the pre-mitoribosome for the association of Mettl15 with Mettl17, we constructed and refined a model of the human pre-mitoribosome with the Mettl15-Mettl17 heterodimer, using the *T. brucei* structure as a template. We started the modeling by superposing human Mettl17 (PDB ID 8CST) and Mettl15 (PDB ID 7PNX) onto the *T. brucei* structure with the conserved rRNA core to obtain a model of the heterodimer. In the initial superposition, a short surface-exposed flexible insertion loop of human Mettl17 (residues 232-240) clashed with Mettl15 at the interface. It exhibits a variable length and sequence among homologs (Figure S2), suggesting it can adopt an alternative conformation compatible with dimer formation. The clash was fixed by relaxation with Amber^47^. Next, we aligned the Mettl15-Mettl17 model onto the early assembly stage structure of the human mitoribosome (PDB ID 8CST) using Mettl17 as an anchor (Figure S3). A knot between residues 304-312 of Mettl15 and residues of 1076-1081 of the rRNA was fixed by rebuilding the protein loop with AlphaFold (see Methods). Here, the position of Mettl15 is rotated by 45° compared to the post-catalytic state. The active site with the cofactor is located more than 40 Å from its target nucleotide C1486. The exact distance could not be calculated, because this rRNA region is disordered in the model. Since the position of Mettl15 is compatible with the human early-stage pre-mitoribosome, we conclude that the modelled intermediate with Mettl15-Mettl17 heterodimer corresponds to a pre-catalytic state (Figure S3).

### Molecular dynamics simulations suggest how Mettl15 recognizes C1486

Since neither reported structures nor our models produced a catalytic state, where Mettl15 would be bound to C1486, we used molecular dynamics simulations to gain insight into the conformational motions that would be required for Mettl15 to reach a catalytically compatible state. Specifically, we asked whether Mettl15-bound SAM is able to closely approach C1486 while Mettl15 maintains its post-catalytic specific interactions with the mtSSU (based on the available post-catalytic state). For this purpose, we used an all-atom structure-based (SMOG^48^) force field, where the post-catalytic structure (PDB ID 7PNX^22^) is explicitly defined to be the global potential energy minimum. Structure-based force fields are well-suited to investigate low-energy motions (i.e. accessible via thermal energy) since they provide predictions of molecular flexibility that are consistent with experimental B-factors^49^ and more detailed explicit-solvent simulations^50^. This has allowed these force fields to be used to characterize molecular flexibility and large-scale conformational rearrangements in mitochondrial and cytosolic ribosomes^4,51^. In the current simulations, we define the rRNA residues proximal to C1483 to be disordered (i.e. residues U1477 to C1494 and A1555 to G1570; see methods), since they are unresolved or have large B-factors in the pre-catalytic state (PDB ID 8CST^26^). Since all non-hydrogen atoms are included in this model, these simulations indicate structural fluctuations that arise from thermal energy are sufficient for Mettl15-bound SAM to closely approach C1486, which is a minimal requirement for catalysis to occur (Figure 4A).

**Figure 4:**
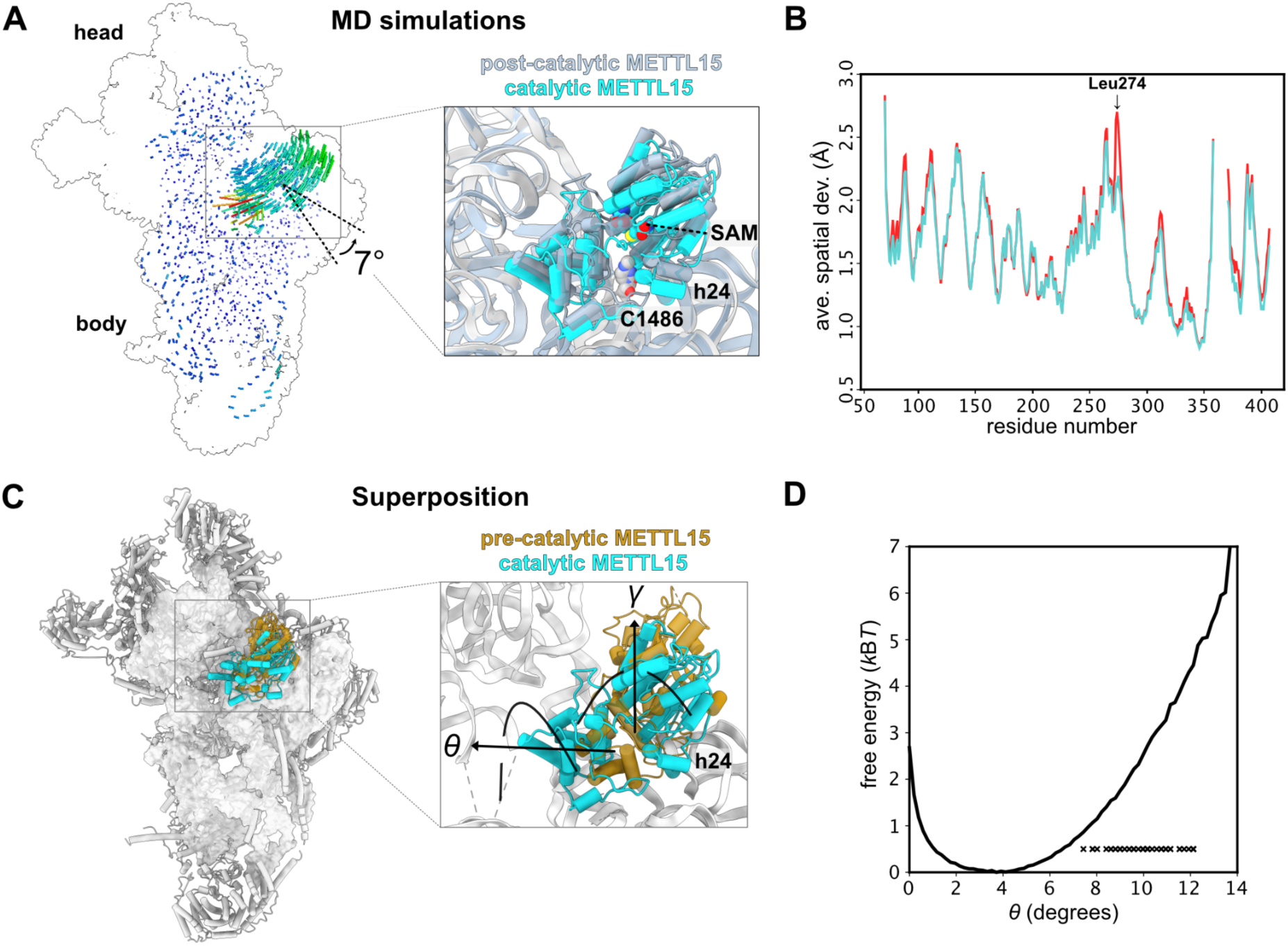
Simulation reveals low-energy structural fluctuations about post-catalytic configuration. **(A)** Comparison of Mettl15 orientations between catalytic (cyan) and post-catalytic (smoky blue) states based on superposition of the mtSSU. Shifts between equivalent Mettl15 Cα atoms and rRNA phosphorus atoms in the different states are color-coded using the spectrum from dark blue to red, corresponding to the range from 0 to 20 Å. **(B)** A.s.d of Mettl15 calculated with respect to the post-catalytic conformation. There are no major intramolecular structural deformations required to adopt orientations in which SAM is proximal to C1486 (labeled “restrained”). The most notable difference is an increase of ∼0.5 Å in Leu274. **(C)** Comparison of Mettl15 orientations between pre-catalytic (brown) and catalytic (cyan) states. The angles describing Mettl15 rotation between the states are indicated. **(D)** Free energy of Mettl15, calculated from a simulation without SAM-C1486 restraints (i.e. unrestrained). When the restraint is included, short distances between SAM and C1486 (<7 Å) can be reached when Mettl15 is rotated/tilted by ∼7-12°. Each “x” indicates a simulated conformation in which the distance is small (<7 Å) in the restrained simulations. These domain orientations are associated with small increases in free energy, indicating that thermal energy is sufficient for Mettl15 to spontaneously adopt catalytically-compatible poses.

Our simulations indicate that large-scale structural deformations in Mettl15 are not required for Mettl15-bound SAM and C1486 to adopt close conformations. To demonstrate this, we introduced a restraint between atom C41 of C1486 and the sulfur atom of the Mettl15-bound SAM molecule. In the simulation, an apparent rotation of Mettl15 is associated with the adoption of short distances (∼6.5 Å) between C1486 and SAM (Figure 4A). These rotated conformations of Mettl15 are associated with small scale bending motions of residues Lys271 to His279. To further characterize these deformations, we performed a second set of simulations in which the restraint was not included. We then compared the average spatial deviation of each residue in Mettl15 after alignment to a post-catalytic structure (Figure 4B). The most significant difference between the restrained and unrestrained simulations was found for residue Leu274, where the average spatial deviation (a.s.d) value increased only slightly, from ∼2.1 Å to ∼2.7 Å.

We also find that short distances between SAM and C1486 can arise through low energy rotational motion of Mettl15. To describe the apparent rotational motion that was present when a restraint was included (above), we calculated the rotation (*γ*) and tilting (*θ*) angles (Figure 4C see methods). The rotation angle (*γ*) was defined as rotation that is parallel to that observed between the pre- and post-catalytic structures. In addition, the tilting angle (*θ*) is defined as rotation that is orthogonal to *γ*. In simulations that included the C1486-SAM restraint, we calculated the rotation and tilt angle for all conformations in which the SAM-C1486 distance was less than 7 Å. This revealed that many of these conformations were associated with low rotation angles (|*γ*| < 3°) and larger tilting angles (7-12°). To probe the energetics of these tilted conformations, we used our unrestrained simulations to calculate the free energy as a function of tilt angle. This showed that tilt angles of 7-12° are only associated with an increase in free energy of ∼ 1-5 k_B_T, relative to the post-catalytic structure (Figure 4D). This indicates that thermally-induced structural fluctuations about a post-like orientation are sufficient for Mettl15 to position SAM within the vicinity of C1486.

### Sequential steps of small mitoribosomal subunit assembly involving Mettl17 and Mettl15

To establish the molecular sequence of Mettl17 and Mettl15 function on the pre-mitoribosome, we ordered the previously obtained structural insights into a series (Figure 5). First, the model from the early assembly stage with TFB1M, along with the *T. brucei*-based model of the Mettl15-Mettl17 heterodimer represents a pre-catalytic state. The next state obtained from molecular dynamics simulations, involves a rearrangement of Mettl15 with a 45° rotation, bringing SAM within 7 Å from the target to provide substrate for its methylation. Since RbfA, and not TFB1M, is present in the model, it is possible that the association of RbfA and the dissociation of TFB1M lead to a disruption of contacts between Mettl15 and Mettl17 resulting in the departure of Mettl17 from the pre-mtSSU. Therefore, only upon the release of Mettl17 can Mettl15 rotate towards C1486 to induce methylation. This sequence of events provides Mettl15 with the conformational space to approach its rRNA target site as predicted by the simulations (Figure 4). Finally, when methylation is accomplished, the conformation of Mettl15 changes again with a backward rotation to adopt a loosely bound state with SAH being 45 Å away from the target. This would ultimately lead to the replacement of Mettl15 by initiation factors in the late stage marking the completion of the mtSSU assembly as previously reported^22^.

**Figure 5:**
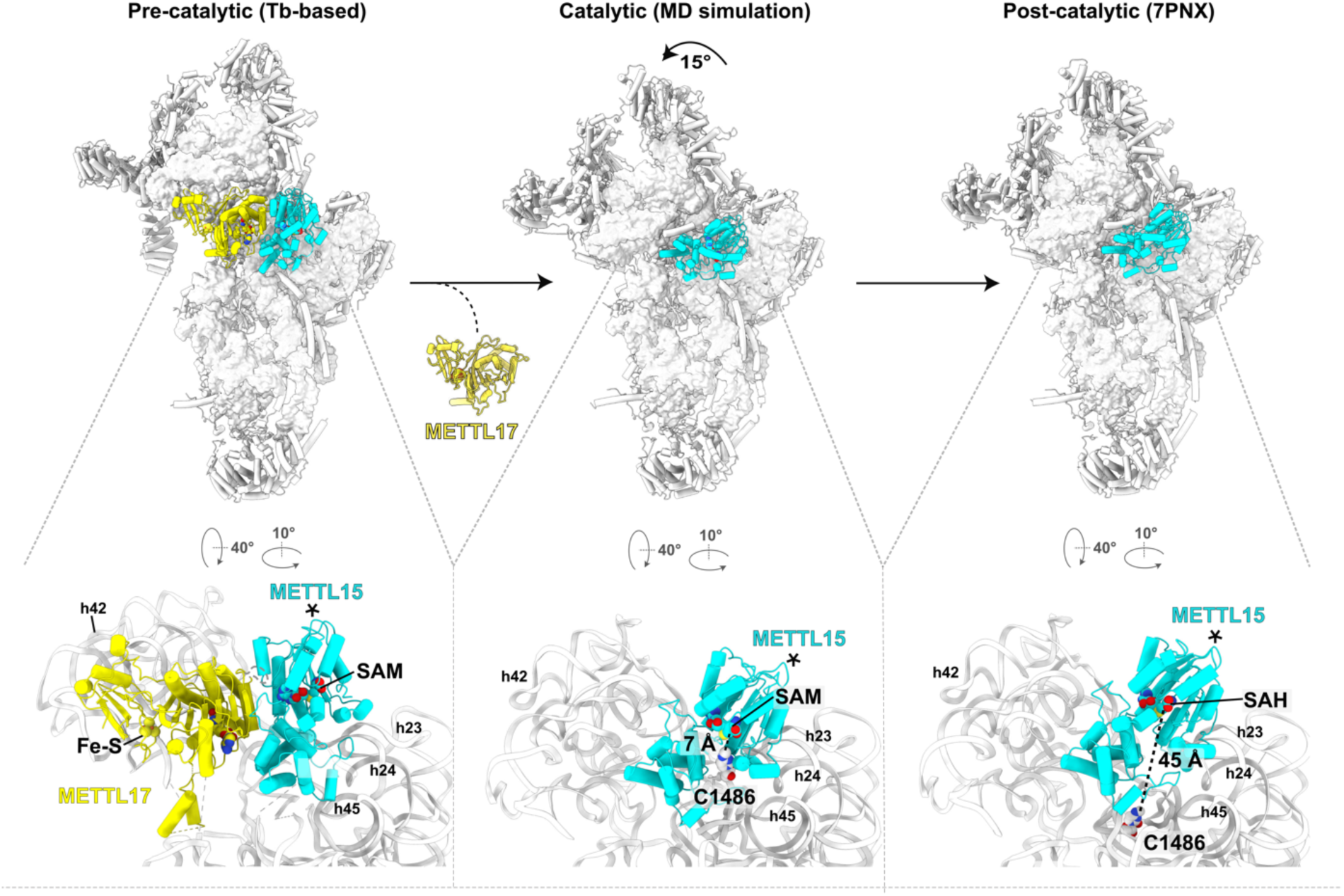
Sequential steps assembly with Mettl17 and Mettl15. Left, *T. brucei* based model of human pre-mtSSU (PDB ID 8CST) with Mettl17-Mettl15. Middle, model from molecular dynamics simulations with rearranged Mettl15 bringing it to the substrate for its methylation. Right, model of the post-catalytic state (PDB ID 7PNX), in which the target nucleotide C1486 is resolved. The estimated distances between the target nucleotide and Mettl15 cofactor SAM are shown and selected rRNA helices are annotated in the close-up views. Asterisks indicate equivalent elements in the three states.

This architecture defines Mettl17 as the key factor that structurally orchestrates the series of assembly events. On one hand, its presence allows the binding of the methyltransferase Mettl15 required for rRNA maturation, and on the other hand, its departure provides the conformational potential of Mettl15 central domains facilitating the rRNA maturation. Thus, Mettl17 stimulates the modification without exhibiting enzymatic activity.

## Discussion

In this analysis, we present *in silico* model of human pre-mitoribosomal assembly, revealing that coupled methyltransferases Mettl15 and Mettl17 are involved in previously undetected, transient assembly states (Figure 6). Our findings indicate that Mettl17 functions as a recruitment factor for Mettl15, forming a structural checkpoint for early assembly stages. This association suggests a broader quality control mechanism where Mettl17, alongside TFB1M, stabilizes Mettl15 and pauses maturation. Release of Mettl17 then facilitates Mettl15’s conformational change on pre-mitoribosome, allowing catalytic methylation of the rRNA, which aligns with observations of the folded rRNA region in this pre-mtSSU assembly^22^. The precise mtSSU head position during C1486 methylation could differ, since it is rotated between pre- and post-catalytic state by 15°. Upon completion of the methylation, Mettl15 is released, and the subunit core can move toward its functional conformation (Figure 6). PyMol^65^ sessions are available for all the states (Supplementary Information). Our integrative structural analysis not only suggests a more complete picture of the mechanistic assembly, but also provides an experimentally-testable hypothesis regarding a potential quality-control mechanism.

**Figure 6:**
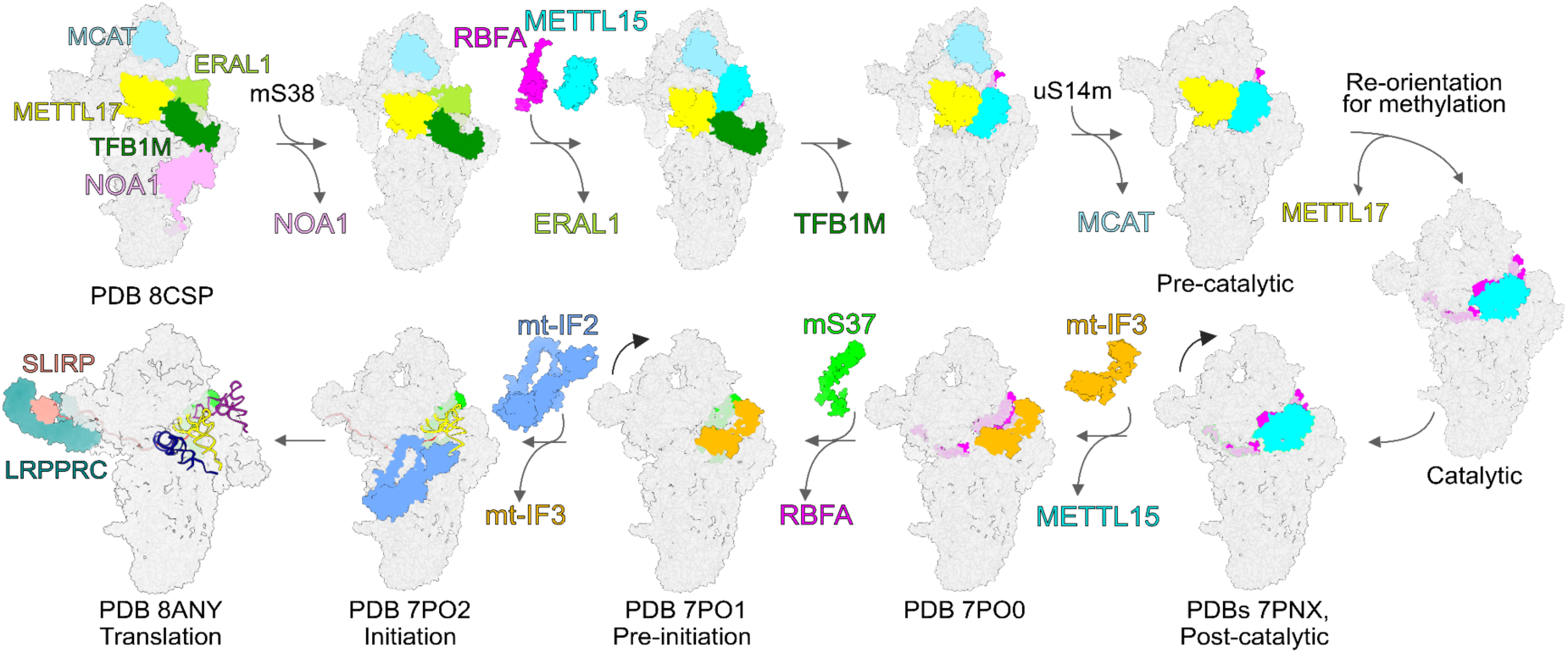
Proposed mtSSU pathway with precursors containing Mettl15-Mettl17 heterodimer and the pre-mtSSU with three methyltransferases. Assembly and initiation factors are shown as colored-coded surfaces. Mitoribosomal proteins and RNA are shown as grey surfaces.

These steps of mitoribosomal assembly are particularly important in the context of biochemical, physiological, and behavioural observations in animals lacking Mettl15^52^. In mice, the loss/ablation/downregulation of Mettl15 has been shown to lead to suboptimal muscle performance, decreased learning capabilities, and lower blood glucose level after physical exercise^52^. The same study, as well as results obtained earlier for cell cultures^36^ also reported accumulation of the RbfA factor, and our model is consistent with these data.

SAM is essential for RNA processing in mouse embryonic fibroblasts and skeletal muscle^53^. Although Mettl17 retains the features typical of class I SAM-dependent methyltransferases^40^, it does not methylate the 12S rRNA region, despite coming into contact with it during assembly. Our models suggesting that Mettl17 acts in recruitment of Mettl15 explain why loss of Mettl17 leads to around 70% reduction in the methylation, resulting in the impaired translation of mitochondrial protein-coding genes and consequent changes in the cellular metabolome^39^. Therefore, it appears that Mettl17 acts as an enhancer of the mtSSU rRNA stability without being directly involved in RNA modification. Targeting of Mettl17 has recently been proposed as a therapeutic approach to suppress colorectal cancer xenograft growth, due to its importance for energy metabolism and maintenance of reactive oxygene species levels during ferroptotic stress^54^. Our findings rationalise the central mechanistic function of Mettl17 in the context of mitochondrial biogenesis. It also provides a more complete description of mtSSU assembly and proposes a plausible explanation for the sequential maturation of the human mitoribosome. Because the two methyltransferases, Mettl15 and Mettl17 co-exist in most eukaryotes, the described functional coupling most likely predates the last common eukaryotic ancestor, and its function presumably became vital as a consequence of the evolution of mitochondrial ribosomes during eukaryogenesis.

Finally, our methodology shows how integrating molecular dynamics with template-based modelling can reveal steps missed in experimental captures due to their transient nature. Although this study has limitations that require further experimental validation, the combined methodology presented here may serve as a more general complementary approach for revealing missing mechanistic steps of transient associations. Together with automated workflows for model building^55,56^, that further integrate diffusion models AF2-predicted structures^57^, scaled up by deep learning systems that generate protein ensembles^58^, this approach can be used for exploring dynamic properties of complex macromolecular systems where only partial experimental data is available. Our work underlines the importance of studying intricate biological processes in combination with advanced computational analyses in order to ultimately predict protein function and derive biogenesis pathways.

## Limitations of the study

In this resorce, we integrate available structural data with model predictions and molecular dynamics simulations to explore previously undetected intermediate states of mtSSU assembly that have eluded experimental characterization. We identified two distinct states and described the features that determine dependence on the specific assembly factor Mettl17 (Figures 5 and 6, Supplementary Information). However, the resulting *in silico* models derived from our analysis require experimental validation. Additionally, we tested only a subset of available structures in this study, which constrained the starting points for our molecular dynamics simulations. As a result, some transitional states may not have been identified using this approach. Moreover, while we propose a plausible composition of the pre-mitoribosome at several stages and identify structural features that dictate the biogenesis pathway, the precise orientations of associated factors and the mtSSU head remain important areas for further investigation. Lastly, although we found that Mettl17 acts as a platform for Mettl15 recruitment and that its subsequent release enables a conformational change in Mettl15 for substrate recognition, understanding the energetic driving forces behind these folding events is an essential next step. Together, future mechanistic experiments in combination with complementary *in silico* approaches presented here is epxeced offer a foundation for elucidating how complex networks of biogenesis and quality-control factors cooperate to ensure the assembly of the catalytic mitoribosome.

**Table S1.** Newly identified features of the *T. brucei* pre-mtSSU.

| **Protein** | **Previous name or chain ID** | **Newly described feature(s)** |
| --- | --- | --- |
| mt-SAF38 | chains UY, Ue | newly identified assembly factor |
| Mettl15 | mt-SAF14 | homolog of Mettl15 (RsmH in *E. coli*)<br>cofactor SAM |
| Mettl17 | mt-SAF1 | homolog of Mettl17<br>cofactor SAM<br>iron-sulphur cluster Fe<sub>4</sub>S<sub>4</sub> |
| RbfA | mt-SAF18 | homolog of RbfA |
| mt-SAF16<br>mt-SAF19<br>mt-SAF25 | - | homologs of *Saccharomyces cerevisiae* Mam33<sup>59</sup> (Uniprot ID P40513), human p32 (Q07021<sup>60</sup>) and *Chlamydomonas reinhardtii* mtSSU protein mS105<sup>61</sup> (A0A2K3DAY3) |
| mS53 | - | residues 63-84 modeled |
| mt-SAF5 | - | residues 560-596 modeled |
| mt-SAF10 | - | residues 4-6 modeled |
| mt-SAF11 | - | residues 148-156 modeled |
| rRNA | - | several regions modeled or adjusted (see Methods) |
| mt-SAF10<br>mt-SAF22<br>mt-SAF27 | chains UB, UC,<br>UD, UF, UI,<br>UJ, UM, UN | ligand acetyl coenzyme A |

## METHODS

### Model building

The PDB ID 6SGB^15^ was used as a starting point and modified as follows. A new assembly factor, mt-SAF38 (Tb927.5.1720), was assigned to an unknown chain in the original model based on the local density (cryo-EM density map EMD-10180) and the presence of the protein in previously isolated trypanosomal mitoribosomal complexes^18^ as revealed by mass spectrometry. Several regions were added or extended in different proteins. In the protein mS53 (chain DF), residues 63-84 were included. N-terminal regions were extended in the models of the proteins mt-SAF10 (chain FA) and mt-SAF11 (chain FB) protein. In the protein mt-SAF5 (chain F5), residues 560-596, previously categorized as an unknown chain, was now modeled. Several proteins have been identified as homologs of assembly factors from other organisms: mt-SAF1 has been assigned as Mettl17, mt-SAF14 as Mettl15, and mt-SAF18 as RbfA. Structural similarity revealed three components of the heterotrimeric assembly mt-SAF16, mt-SAF19, and mt-SAF25 are homologs of the homotrimer-forming human protein p32, yeast Mam33, or algal mS105.

The model of rRNA was modified as follows. The linker between nucleotides 560-620 (h44, h45) was adjusted. Some regions with insufficient resolution quality of density were removed, namely nucleotides 208-226, 254-260, 349-353, 385-389, 397-417, 431-440, 489-510, and 523-529. Nucleotides 67-73, 80-87, 171-172, 183-189, 273-285, 322-324, and 366-374 were shown as ribose-phosphate backbones.

Several ligands were included in the model. Consistent with previous observations^23^, we identified the density corresponding to GTP in the protein mS29 and PO_4_^3-^ in the protein mt-SAF29. Ligand in Mettl17 and Mettl15, originally assigned as S-adenosylhomocysteine (SAH) molecules were substituted with S-adenosylmethionines (SAM), because there is no evidence suggesting that these proteins exist in the post-catalytic state, and the presence of SAM is more plausible in the context of our results. Furthermore, the Zn^2+^ ion present in Mettl17 was replaced with an Fe_4_S_4_ iron-sulfur cluster, consistently with the density, identity of coordinating residues and recent identification of iron-sulfur cluster in yeast and mammalian homologs. Unidentified chain UD was replaced with acetyl Co-A.

### Structure prediction and analyses

Structure prediction was performed by AlphaFold3^30^ or AlphaFold Multimer^28^. The latter was used with databases BFD, Mgnify 2018_12, UniRef30 2021_03, UniRef90 2023_04 to predict structures and calculate ipTM scores for Mettl15 in dimer with all assembly factors present in early mtSSU intermediate. For *T. brucei* homology model, the clash at the interface between Mettl15 and Mettl17 was fixed by running a relaxation procedure with the Amber potential as done in AlphaFold, and the knot between Mettl15 and RNA was fixed by inpainting as described in AF_unmasked^64^. Here, the Mettl15-Mettl17 complex was used as a multimeric template and the clashing loop was deleted from the template so that it could be rebuilt by AlphaFold. Fifty predictions were generated this way, and the one closest to the initial template (RMSD: 0.2) where the clash would be fixed when including the RNA was selected. We neither show nor interpret regions with pLDDT scores below 65 in any of the models. Angles between Mettl15 in different models were calculated using the PyMOL (Schrödinger, US)^65^ built-in script angle_between_domains.

### Identification and phylogenetic analyses of Mettl15 and Mettl17 across eukaryotes

Using *Escherichia coli, Homo sapiens*, and *Trypanosoma brucei* orthologs of Mettl15 and Mettl17 as queries for blastp search against the EukProt v3 database^66^, we built starting datasets that were subsequently cleaned from apparent eukaryotic contaminations using phylogenetically-aware approach (identification of possible contaminants by visual inspection of phylogenetic tree followed by manual check of their origin). Cleaned datasets were used to build profiles hmm in HMMER3^67^. Next, 131 organisms that cover known eukaryotic diversity and whose genome or transcriptome assemblies are of a good quality were selected for the final search. This search was performed in three steps: 1/ HMMER3 search with profiles hmm; 2/ blastp search^68^ using query sequence from a closely related species; 3/ tblastn search in corresponding nucleotide assembly (to exclude possibility that ortholog is missing due to an inaccurate protein prediction). Names of selected organisms, accession numbers of used assemblies, and tools that were used for successful search are indicated in the Table S2. Multiple sequence alignments of the homologous amino acid sequences were built using MAFFT v7.407 with the L-INS-i algorithm^69^ and were manually trimmed to exclude unreliably aligned regions. The maximum likelihood tree was inferred with IQ-TREE multicore v2.2.0.3^70^ using the LG4X substitution model. Statistical support was assessed with 100 IQ-TREE non-parametric bootstrap replicates. Sequences of both genes from all organisms are available in Supplementary Data 1&2.

### Molecular dynamics simulations

#### Potential energy function

An all-atom structure-based “SMOG” model^48^ of the mitoribosome small subunit was used to probe the scale of structural fluctuations around the post-catalytic state and determine whether thermal energy is sufficient for Mettl15-bound SAM to approach C1486, or whether Mettl15 is more likely to be associated with a larger-scale rearrangement that would require transient dissociation from the ribosome. The force field that was used is a single-basin model where the post-catalytic structure (PDB ID 7PNX) was defined as the global potential energy minimum. The specific variant of the force field is available through the smog-server force field repository (https://smog-server.org), with entry name AA_PTM_Hassan21.v2. The functional form of the potential energy is given as:

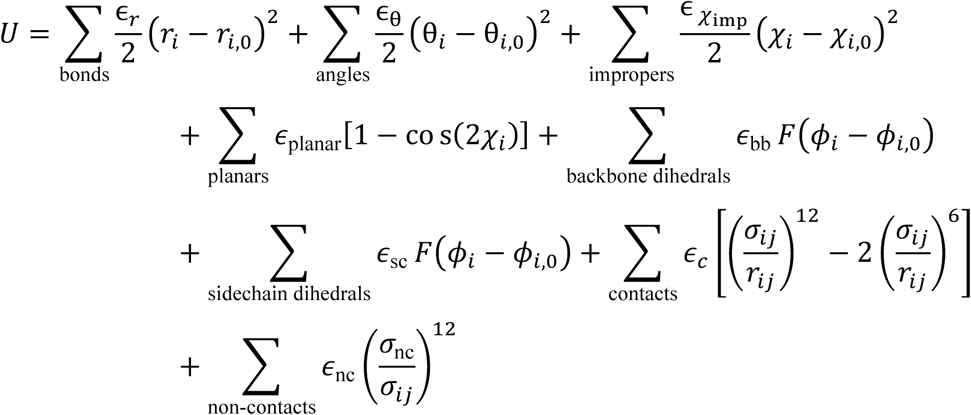

where

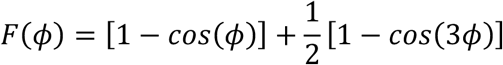

{*r*_0_} and {*θ*_0_} parameters are given values found in the Amber ff03 force field^71^. Dihedral parameters {*χ*_0_} and {*ϕ*_0_} are assigned the corresponding values found in the experimental model. Non-bonded contacts that are found in the experimental model, are identified according to the Shadow Contact Map algorithm, with a shadowing radius of 1 Å and a cutoff distance of 6 Å. The contacts are given an attractive 6-12 interaction that stabilizes the preassigned structure, with interatomic distance *σ*_*ij*_ that is found in the experimental structure, multiplied by 0.96 to avoid artificial expansion of the structure^72-74^. Atom pairs that are not in contact are assigned a repulsive potential to model excluded-volume steric interactions, where *σ*_*nc*_ is given the value 2.5 Å. Energy scale weights are defined as 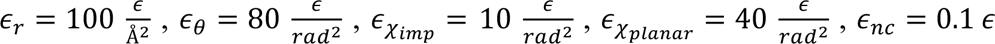, where *ϵ* is the reduced energy unit. The dihedral and contact energy weights are normalized as in Whitford et al, Proteins 2009.

Since rRNA residues near the Mettl15 binding site are disordered (unresolved or high B-factors) in the pre-catalytic structure, these regions were modelled as disordered. For this, stabilizing contacts and dihedrals for the flexible rRNA region (i.e. residues U1477 to C1494 and A1555 to G1570), were removed.

Two sets of simulations were performed. In the first set of simulations, a harmonic restraint was introduced, which ensured that the distance between C1486 and SAM (atom name) adopted short values. The harmonic restraint had a minimum at 5 Å and the spring constant was 150 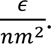 These simulations were used to ask whether simple bending motions of Mettl5 are sufficient for SAM and C1486 to become proximal. In the second set of simulations, the restraint was not included. These unrestrained simulations were performed to determine the scale and direction of structural fluctuations that can arise from thermal energy.

#### Simulation details

All force field files were generated using SMOG2 software package^48^. Molecular dynamics simulations were performed using OpenMM^74^ and OpenSMOG^73^ libraries. The simulations were performed at a reduced temperature of 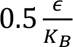 that was maintained by using Langevin dynamics protocols.

#### Calculating rotation angles for Mettl15

Euler angles were used to describe rotation of Mettl15, relative to the mtSSU body^71,72^. Consistent with methods for describing rotation of the mtSSU^71^, we described a rotation angle *γ*, which is the sum of the *ψ* and *ϕ* angles in the Euler formulation (Figure S4). The polar angle *θ* (i.e. tilt angle) represents rotation that is orthogonal to the primary rotation. To calculate Euler angles, we first assigned a set of axes that remain fixed in the frame of reference of Mettl15. For convenience, we define the “Z” axis as the axis of rotation (Euler-Rodrigues axis) between the pre and post catalytic states. The following protocol was used to define the primary rotation axis:

1. Least squares alignment of the mtSSU (excluding Mettl15) of the pre-catalytic structure to post-catalytic structure.
2. Associate coordinate system to Mettl15 of pre and post catalytic structures.
3. The E-R angle was then calculated between the coordinate systems of both structures. The angle was found to be 45°.

To calculate Euler angles in the simulations, the following protocol was applied:

1. Define “Z” axis as the E-R axis. This ensures that our primary rotation angle *γ* describes rotation that is parallel to that defined by the pre-to-post rearrangement.
2. Align each simulated frame to the post-catalytic structure of the mtSSU, where alignment was based on non-Mettl15 atoms.
3. Align the post-catalytic conformation of Mettl15 to each simulated frame.
4. Calculate the Euler angles (*ϕ*, 𝜓 and *θ*) between the post-catalytic and aligned (previous step) orientation.
5. Define rotation as *γ* = *ϕ* + 𝜓.
6. Define tilt as *θ*.

### Source data

The atomic model of the *T. brucei* mtSSU precursor was deposited in the PDB database (PDB ID 9HNY). All data from phylogenetic and structural analyses are available as supplementary material or have been deposited on Figshare (link will be provided in the accepted version).

## Acknowledgements

This work was supported by the European Research Council (ERC-2018-StG-805230), Czech Science Foundation (20-04150Y) to O.G., the project P JAC CZ.02.01.01/00/22_008/0004575 RNA for therapy, co-funded by the European Union, and the Ministry of Education, Youth and Sports of the Czech Republic through the e-INFRA CZ (ID:90254) to O.G. and A.Z., and SciLifeLab BeyondFold to B.N. G.W. and P.C.W were supported by NIH grant R35GM153502-01. Some of the structure prediction experiments and other analyses were enabled by the Berzelius resource provided by the Knut and Alice Wallenberg Foundation at the National Supercomputer Centre in Sweden. Work in the Center for Theoretical Biological Physics was supported by the National Science Foundation (NSF) grant PHY-2210291. We thank members of Amunts lab for their contributions to model building, data interpretation, and discussions.

## Author contributions

Y.Z. built the model; C.M. carried out computational modeling and structure prediction; G.W. and P.C.W. performed molecular simulations; Y.Z., C.M., G.W., P.C., P.C.W., O.G., A.A. performed the structural analysis; T.P. performed the phylogenetic analysis; B.N., A.Z. supervised the project; A.A. and O.G. wrote the manuscript with help from Y.Z., P.C.W. All the authors contributed to the manuscript preparation.

---

**Figure S1.**
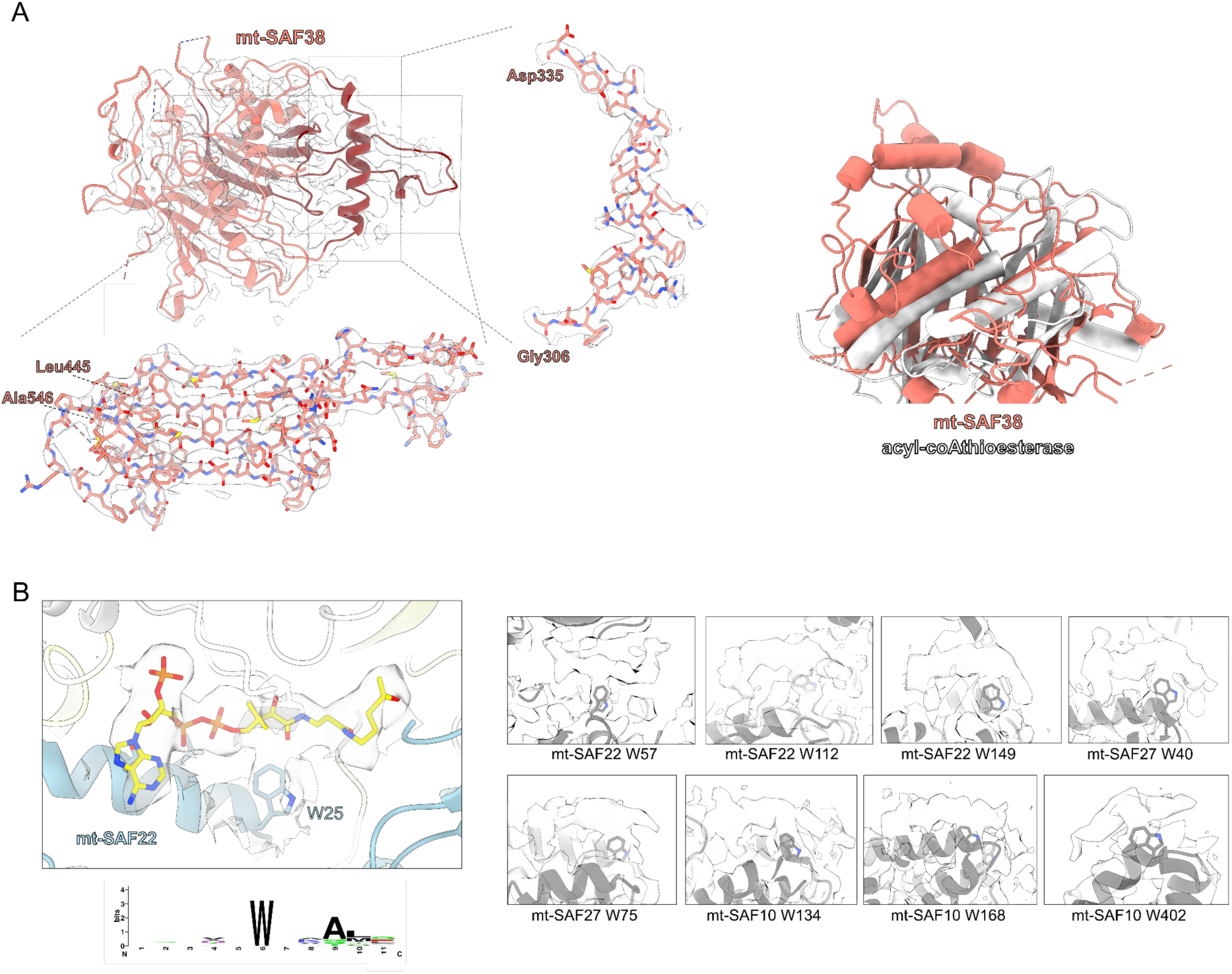
New features in the map. **(A)** The density and model of the newly identified mt-SAF38 with close-up views showing how residues fit the density. Right, mt-SAF38 (dark red) superposed with mouse acyl-coA thioesterase (PDB 5ZV3). **(B)** Acetyl-CoA placed into a density associated with tryptophan 25 of mt-SAF22 and other examples of the hammerhead shaped densities. Sequence logo of acetyl-CoA binding regions, showing the conserved tryptophan, was created using WebLogo^75^.

**Figure S2:**
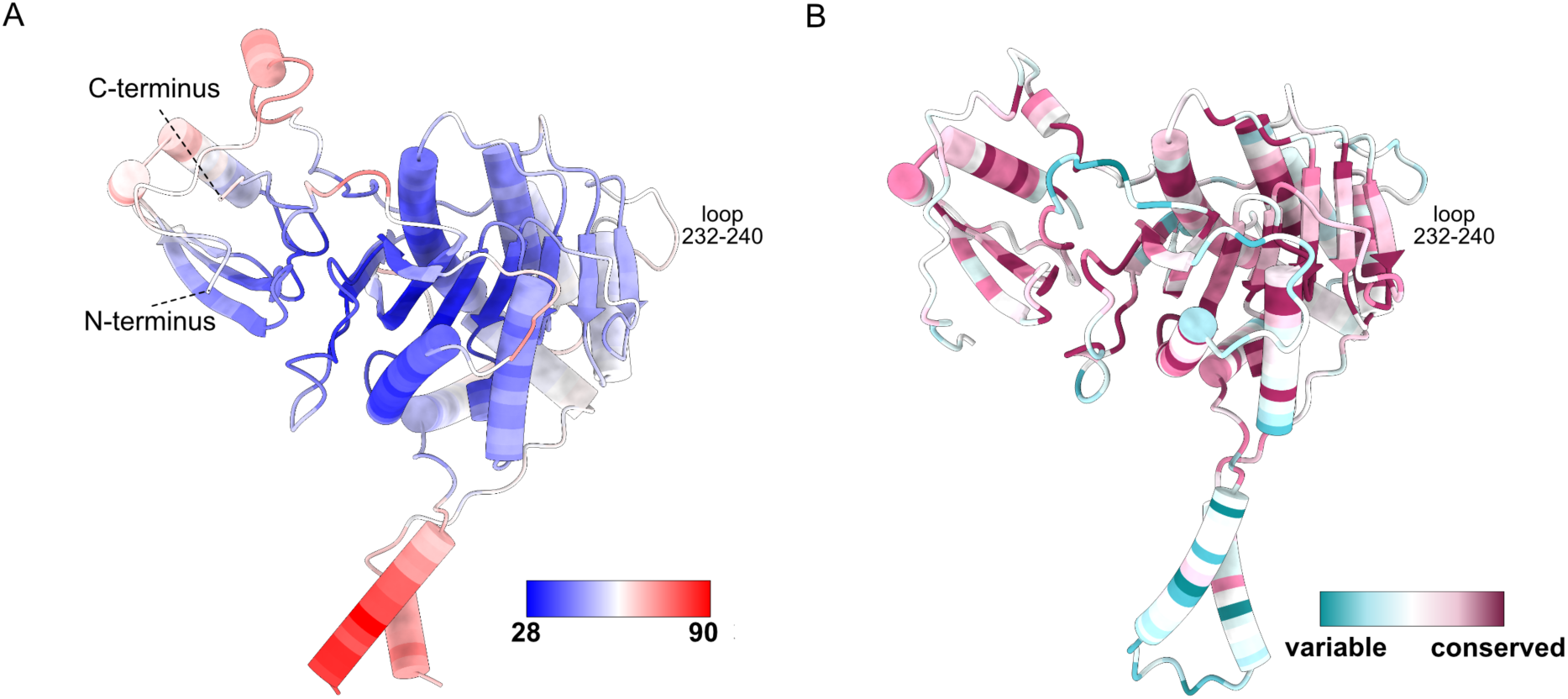
Flexibility and conservation of Mettl17. **(A)** PDB ID 8CST colored by B-factor. Increased flexibility of the loop 232-240 is evident by a higher B-factor. **(B)** The conservation coloring profile calculated by ConSurf repository^76^ mapped onto the model.

**Figure S3:**
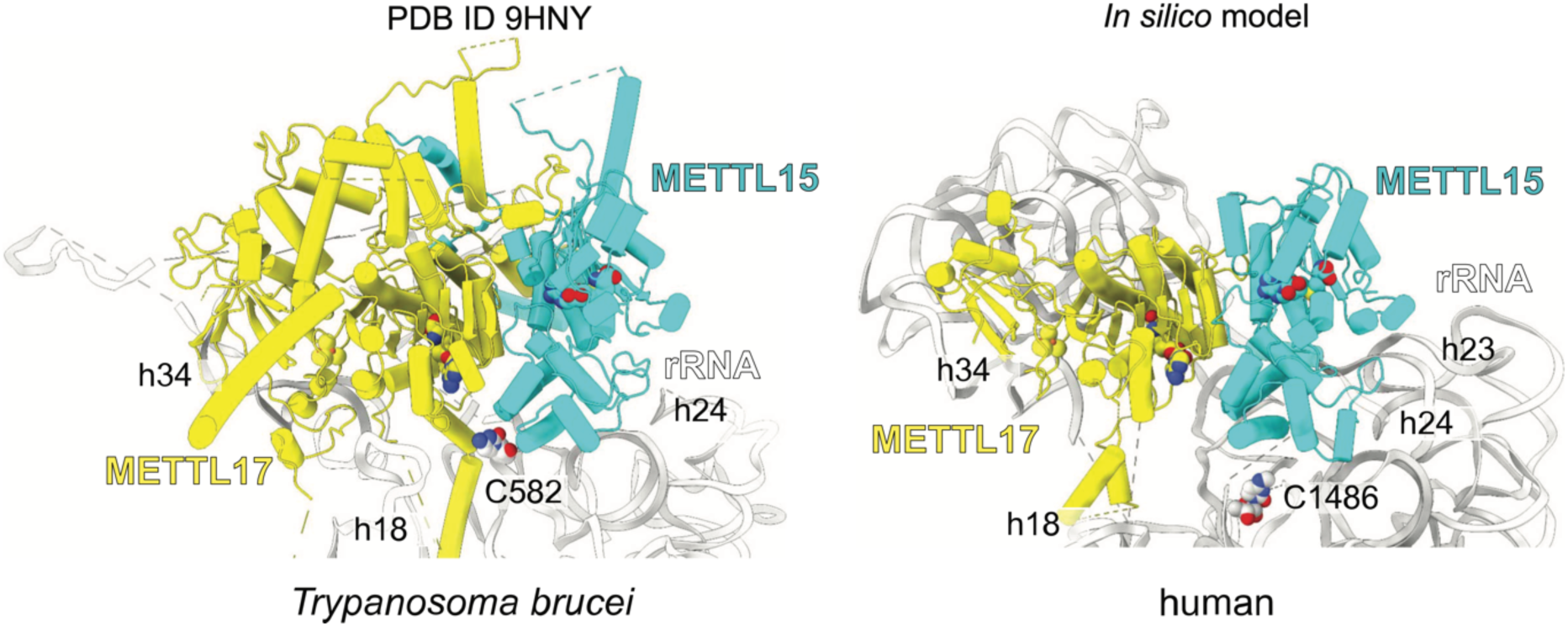
Comparison of the Mettl15-Mettl17 heterodimer in *T. brucei* and corresponding *in silico* model of the human mitoribosome. Human early-stage pre-mitoribosomal model PDB ID 8CST was aligned onto *T. brucei* Mettl17, and Mettl15 was modelled based on the trypanosomal template with no clashes. The position of Mettl15 in the created *in silico* model is compatible with the experimental data.

**Figure S4:**
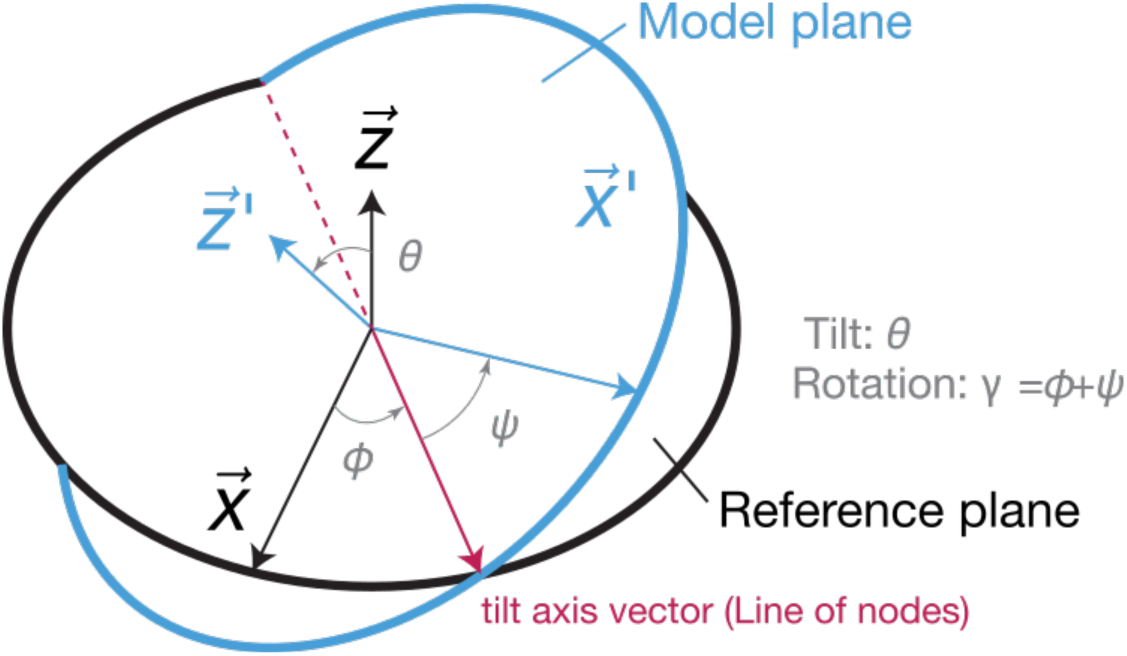
Description of Euler angles used to analyze molecular dynamics simulation. The angles were calculated by comparing the vectors of the model plane *z*^⃑′^, *x*^⃑′^ with the corresponding vectors of the reference plane *z*^⃑^, *x*^⃑^. Rotation is defined be angle *γ* = *ϕ* + 𝜓, while tilt is defined by angle *θ* around the tilting axis (line of nodes).

**Resource: PyMOL Session File.** Aligned structures of 11 states, each with annotated assembly factors, rRNA, and mitoribosomal proteins, coloured as in Figure 6.

